# Integration of anatomy ontology data with protein-protein interaction networks improves the candidate gene prediction accuracy for anatomical entities

**DOI:** 10.1101/2020.03.07.981795

**Authors:** Pasan Chinthana Fernando, Paula M Mabee, Erliang Zeng

**Affiliations:** Department of Biology, University of South Dakota, Vermillion, SD, USA; Division of Biostatistics and Computational Biology, College of Dentistry, University of Iowa, Iowa City, IA, USA; Department of Preventive & Community Dentistry, College of Dentistry, University of Iowa, Iowa City, IA, USA; Department of Biostatistics, College of Public Health, University of Iowa, Iowa City, IA, USA; Department of Biomedical Engineering, College of Engineering, University of Iowa, Iowa City, IA, USA

**Keywords:** Anatomy Ontology, Phenotype, Protein-protein interaction networks, Big data, Data integration, Candidate gene prediction, Data quality, Semantic similarity

## Abstract

**Background:** Identification of genes responsible for anatomical entities is a major requirement in many fields including developmental biology, medicine, and agriculture. Current wet-lab techniques used for this purpose, such as gene knockout, are high in resource and time consumption. Protein-protein interaction (PPI) networks are frequently used to predict disease genes for humans and gene candidates for molecular functions, but they are rarely used to predict genes for anatomical entities. This is because PPI networks suffer from network quality issues, which can be a limitation for their usage in predicting candidate genes for anatomical entities. We developed an integrative framework to predict candidate genes for anatomical entities by combining existing experimental knowledge about gene-anatomy relationships with PPI networks using anatomy ontology annotations. We expected this integration to improve the quality of the PPI networks and be better optimized to predict candidate genes for anatomical entities. We used existing Uberon anatomy entity annotations for zebrafish and mouse genes to construct gene networks by calculating semantic similarity between the genes. These ‘anatomy-based gene networks’ are semantic networks, as they are constructed based on the Uberon anatomy ontology annotations that are obtained from the experimental data in the literature. We integrated these anatomy-based gene networks with mouse and zebrafish PPI networks retrieved from the STRING database, and we compared the performance of their network-based candidate gene predictions.

**Results:** According to candidate gene prediction performance evaluations tested under four different semantic similarity calculation methods (Lin, Resnik, Schlicker, and Wang), the integrated networks showed better receiver operating characteristic (ROC) and precision-recall curve performances than PPI networks for both zebrafish and mouse.

**Conclusion:** Integration of existing experimental knowledge about gene-anatomical entity relationships with PPI networks *via* anatomy ontology improves the network quality, which makes them better optimized for predicting candidate genes for anatomical entities.

## Background

Unraveling molecular and phenotypic functions of proteins is a cornerstone in molecular biology. In particular, understanding the genes associated with the formation of anatomical structures, also termed ‘anatomical entities’, is essential in developmental biology [1–4]. The majority of genes associated with anatomical entities are obtained using wet-lab methods, such as gene knockout [5, 6], gene knockdown [7], and overexpression [8, 9]. These methods, however, are time-consuming and require significant resources, and thus only a few genes may be associated with the development of a particular anatomical entity, though there are likely many more genes involved.

Alternatively, computational prediction methods for discovering gene-anatomical entity associations can be employed because of their higher speed and low resource consumption. Sequence similarity-based function prediction is such an example, which is widely used to predict the molecular functions of proteins [10, 11]. However, using it to predict anatomical associations of genes is questionable, because anatomical entities develop from a combination of several biological pathways that include proteins with diverse molecular functions and sequences [12]. On the other hand, protein-protein interaction (PPI) networks can be used to predict candidate genes for anatomical entities, based on the assumption that proteins that regulate the same term or function are more likely to physically interact with each other [13, 14]. PPI networks represent such interactions as graphs where proteins are represented by nodes and their interactions are represented by edges. PPI networks have been widely used in predicting candidate genes for human disease phenotypes [15–17]. Therefore, PPI networks are suitable for predicting candidate genes associated with anatomical entities. However, the challenge with PPI network-based candidate gene prediction is improving the accuracy of the predictions [13, 18–21], which is low because of the poor quality of the large-scale PPI network data sets [14, 21–23]. PPI networks are generated by experimental methods such as yeast two-hybrid assay and high-throughput mass-spectrometric protein complex identification (HMS-PCI), which can generate false positive interactions [19]. Furthermore, PPI networks for model organisms are still incomplete and the quality of data varies depending on the model [18, 24]. For instance, well studied organisms such as human and mouse contain more complete PPI network data sets compared to *Xenopus* or zebrafish [14, 20].

The STRING database is the most widely used PPI database, and it currently contains PPI networks for 2031 organisms [18, 20, 25, 26]. To improve the quality of PPI network data, the STRING database also computationally predicts the strength of an interaction between two proteins based on properties such as co-expression in addition to experimental evidence. This results in additional quality-controlled PPI network datasets. Although STRING database does not incorporate experimental evidence regarding proteins that are regulating similar anatomical entities, we assume two proteins that regulate similar anatomical entities are more likely to interact with each other.

The information regarding gene-anatomical entity associations that are discovered *via* wet lab techniques such as gene knockout is recorded in literature, annotated to model organism databases using gene and anatomy ontologies, and is available through these databases and integrated repositories such as the Monarch Initiative [27, 28]. The anatomical entity associations of genes in the Monarch Initiative repository are annotated using Uberon anatomy ontology entities [29–31]. Such ontology annotations enable calculation of the functional similarity between any two genes using computational methods [32–35]. By calculating the pairwise similarity for all gene pairs with anatomical annotations, a semantic gene network can be constructed based on genes’ anatomical annotations. Such a network is referred to as an ‘anatomy-based gene network’. The concept of constructing gene networks using ontology information has been previously presented [17, 36–38], but it was applied only using the Gene Ontology (GO). For example, Jiang, et al. [37] constructed a gene network using the Gene Ontology-Biological Process (GO-BP) entities to infer disease genes in humans. This gene network, however, was not integrated with an existing PPI network; instead, it was used directly for disease gene prediction and the results were compared with a human PPI network. They discovered that the semantic gene network that uses GO outperformed the PPI network when predicting disease genes [37]. Zeng, et al. [36] performed gene function prediction using both PPI data and GO annotations, where the semantic similarity between GO terms was used to derive semantic similarity between genes, which was then used to evaluate the quality of PPI data [36]. This is based on the conclusion that semantic similarity between genes serves as a metric of support for PPI data [39], which even can be used to predict linkages between genes to generate a functional gene network [40].

Motivated by these results, we here integrate known gene-anatomical entity associations retrieved from the Monarch Initiative repository with STRING PPI networks for zebrafish and mouse in an attempt to improve the quality of the networks and the prediction accuracy of novel gene candidates for anatomical entities, such as limbs and fins. Here, the focus is to construct semantic similarity gene networks based on anatomy ontology annotations (anatomy-based gene networks) using multiple semantic similarity methods and integrate them with the PPI networks retrieved from the STRING database for the same species (zebrafish and mouse). This method assigns higher weights to the gene interactions that are supported by both source networks; we expect this to enhance the quality of the source networks by reducing the false positive and false negative interactions (Figure 1).

**Fig. 1.**
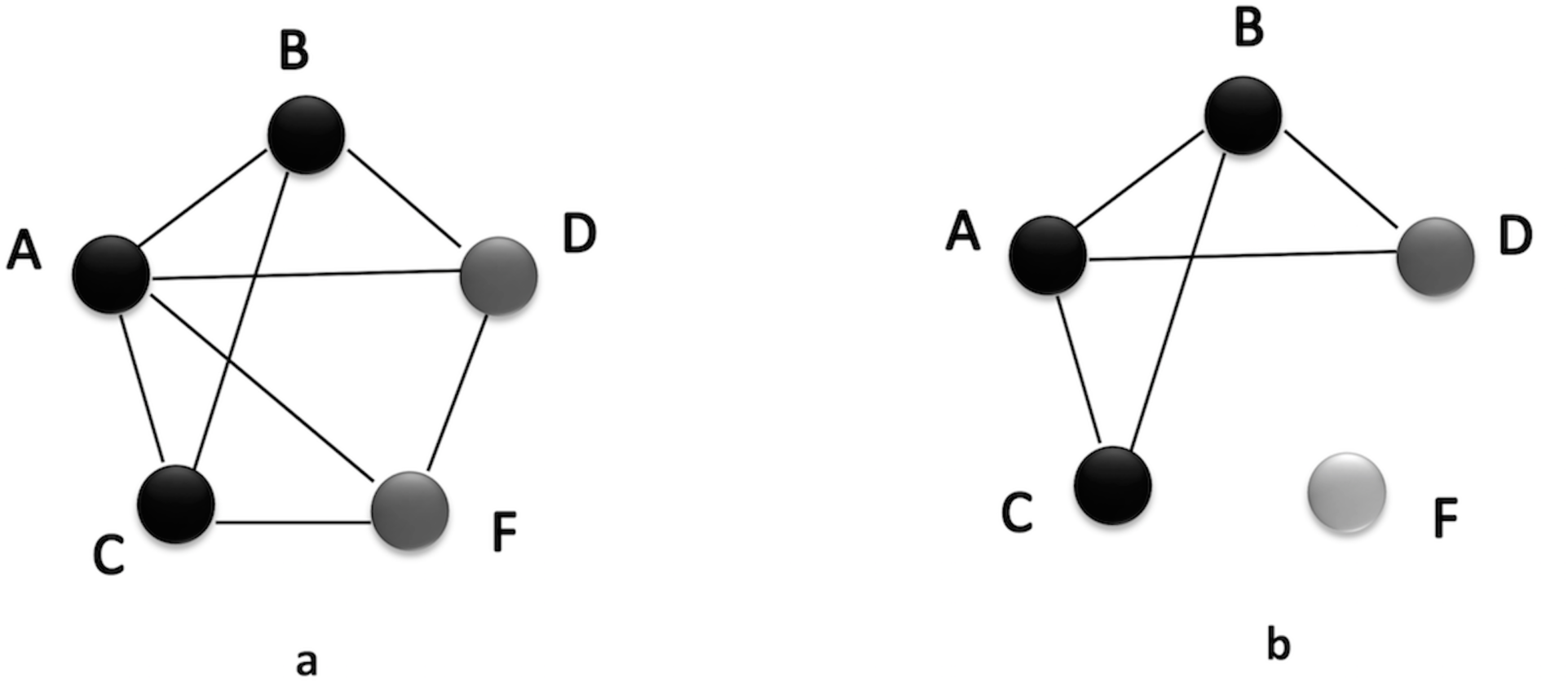
A hypothetical scenario that compares candidate gene predictions based on a (a) PPI network and an (b) anatomy-based gene network. The nodes A, B, and C (colored in black) in both networks represent three genes known to be associated with a certain anatomical entity denoted as *entity 1*. In the PPI network (a), genes D and F are predicted to be associated with *entity 1* because genes D and F interact with genes A, B, and C that are known to be associated with *entity 1*. In contrast, the anatomy-based gene network (b) only predicts D as a potential candidate for *entity 1* because the gene F does not have any interaction with other genes annotated with *entity 1*. The absence of interactions of gene F in gene network (b) can be due to two reasons: (1) it is not annotated with any anatomical entities, (2) it is not annotated with entities that are similar to the anatomy entities associated with genes A, B, or C. The anatomy-based gene network (b) is built entirely on anatomy ontology information, thus it provides a different interaction structure. Hypothetically, the gene F could be a false positive interaction in the PPI network, and the integrative use of the anatomy-based gene network may reduce the false positives by filtering them.

## Methods

### Data sources

PPI networks for zebrafish and mouse were retrieved from the STRING database [41]. The proteins in the PPI networks were represented by unique STRING ID, and we replaced them with corresponding gene names/symbols (retrieved from STRING database) to facilitate network integration in later stages. Usually, raw PPI networks are too large for downstream analyses. Therefore, we filtered the networks based on the recommended 0.7 gene interaction/combined score cutoff [18].

To construct anatomy-based gene networks, initially, anatomical profiles must be constructed. An anatomical profile represents the multiple anatomical entity annotations for a gene. We obtained known gene-anatomical entity relationships for zebrafish and mouse from the Monarch Initiative repository [27, 28]. The Monarch Initiative retrieves genes and their anatomical entity annotations for zebrafish and mouse from the zebrafish [42] and mouse [43] model organism databases, respectively, and associates them with the corresponding Uberon anatomy entities. The annotations available in the Monarch Initiative are pre-processed and cross-checked with other model organism annotations to remove uncertain gene-anatomical entity associations that result when the expression of multiple genes are simultaneously disrupted to observe the effect on a given phenotype [27].

We arranged the gene-anatomical entity associations into the following format where *G_1_* and *G_2_* represent two genes, and *(t_a1_, t_a2_ … t_am_)* and *(t_b1_, t_b2_ … _tbn_)* represent their associated anatomical entities (Uberon entities), respectively.

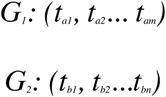

Gene name/symbol reconciliation was required to align data from STRING to Monarch. The STRING database obtains data from various data sources, such as Entrez Gene database [44] and UniProt knowledgebase [45], whereas the Monarch Initiative data repository obtains data from model organism databases. Occasionally, the gene names do not match. Therefore, we computationally matched gene names of the two data sources using three steps: (1) match the genes directly using their names/symbols, (2) match using their Ensembl identifiers, and (3) match the remaining gene names in the anatomical profiles to the synonyms available in the STRING database. Each step attempted to sequentially minimize the number of gene mismatches.

### Construction of the anatomy-based gene networks

To construct anatomy-based gene networks, we first calculated semantic similarity scores between Uberon anatomy entities annotated to gene pairs, and then, aggregated those scores to calculate the gene similarity scores between all gene pairs (Figure 2).

**Fig. 2.**
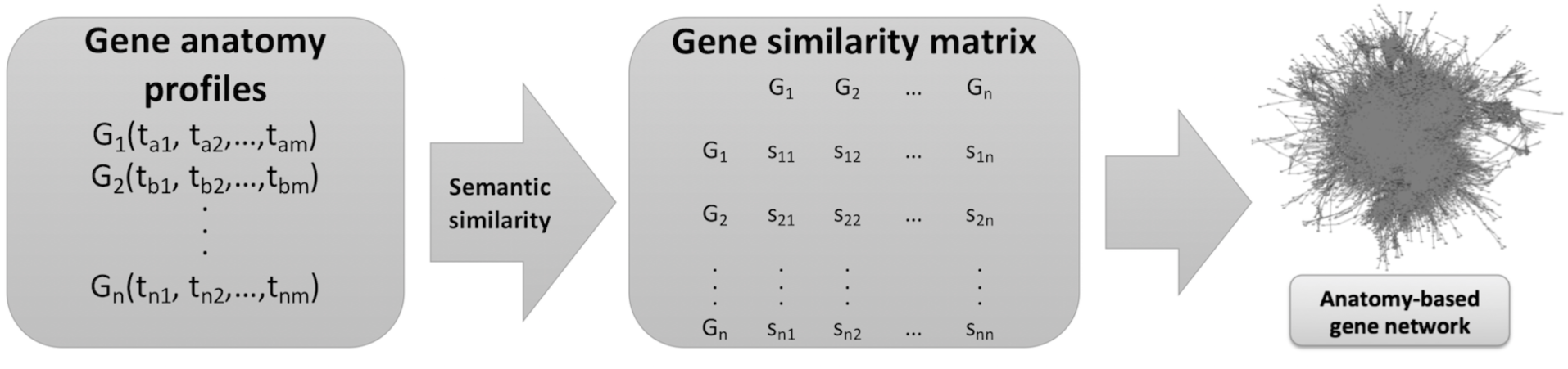
The general workflow for generating anatomy-based gene networks. The genes are represented by *G*_1_, *G*_2_, etc., and their anatomical entities (Uberon terms) are represented by *t_a1_*, *t_b1_*, etc. In the gene similarity matrix, the similarity scores between genes are represented by *s_11_*, *s_12_*, etc.

We used four semantic similarity methods to calculate semantic similarity between Uberon terms: Wang method [46], Resnik method [47], Lin method [48], and Schlicker method [49]. The latter three methods are based on calculating the information content (IC) of each term in the ontology hierarchy, and their equations are given below (equations 1, 2, and 3).

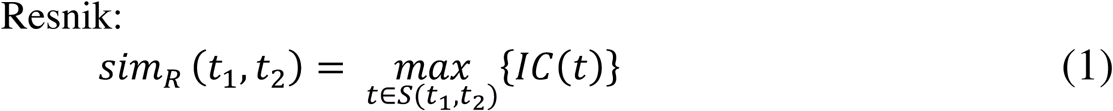

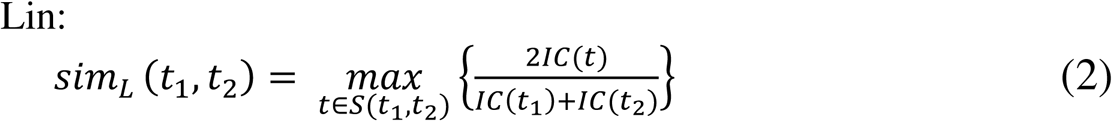

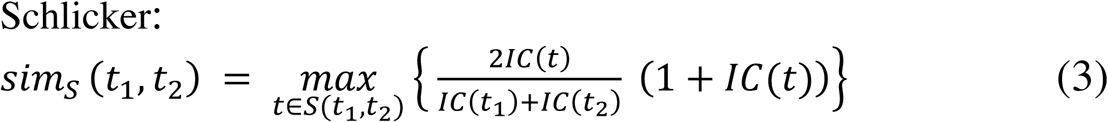

In the above equations, *t_1_* and *t_2_* represent the ontology terms between which the similarity is calculated, whereas *S* denotes the set of common ancestors for the two terms. The information content for a given term *t* is represented by *IC(t)*, which is calculated based on the number of genes annotated to the term *t* as illustrated below (equations 4 and 5).

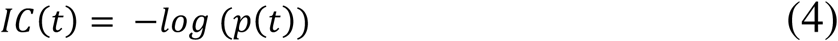

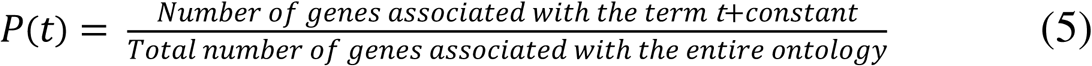

The Wang method does not use the information content [46]. It only depends on the ontology structure and the relationships between the entities. The equation for the Wang semantic similarity calculation between the two entities *t_1_* and *t_2_* is given below.

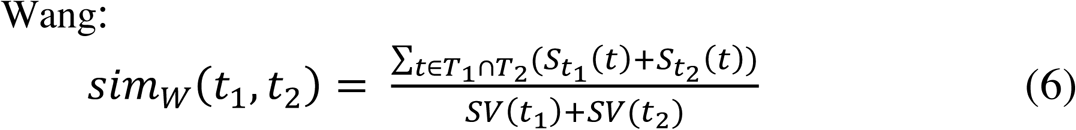

In the above equation, *S_t__i_*(*t*) represents the semantic contribution of term *‘t’* on term *t_i_*, when *‘t’* is an ancestor of *t_i_*. The term *SV*(*t_i_*) represents the semantic contribution of all the ancestors of term *t_i_* on itself.

We used the above four semantic similarity calculation methods to calculate the similarity between gene pairs using the method explained below. For instance, if the gene *G*_1_ is annotated with the Uberon entities: (*t_a1_, t_a2_ … t_am_*), and the gene *G*_2_ is annotated with the Uberon entities: (*t_b1_, t_b2_ … t_bn_*), then the similarity between the two genes, *sim* (*G_1_, G_2_*), can be calculated using the following equation 7.

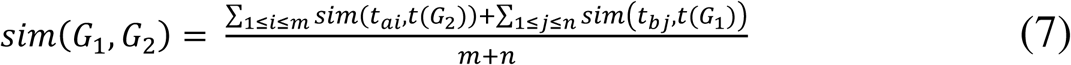

The *t*(*G*_1_) = (*t_a1_, t_a2_ … t_am_*) *and t*(*G*_2_) = (*t_b1_, t_b2_ … t_bn_*) represent Uberon anatomy profiles for gene *G*_1_ and gene *G*_2_, respectively, and the *sim*(*t_ai_, t*(*G*_2_) represents the maximum semantic similarity between term *t_ai_* and any of the entities in *t*(*G*_2_), which can be calculated using the equation 8 below.

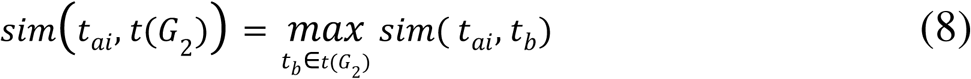

Using the equation 7, we calculated a similarity score for each gene pair in the anatomical profiles for zebrafish and mouse, which generated a pairwise gene similarity matrix for each semantic similarity calculation method.

After obtaining a gene similarity matrix, the final step of the anatomy-based gene network construction was to connect each pair of genes with an edge. This can be done with or without applying a gene similarity cutoff. Without the cutoff, all the gene pairs with a similarity score are retained in the anatomy-based gene network (unfiltered network). With the cutoff, if the pairwise gene similarity score between two genes is higher than the cutoff, an edge will be placed to connect the two genes, otherwise, they will not be connected (filtered network). We obtained suitable cutoffs for anatomy-based gene networks by analyzing their similarity score distributions, which were selected to keep the number of interactions/gene pairs approximately similar to that of the STRING PPI networks with the 0.7 cutoff for each model organism.

### Integration of the anatomy-based gene networks with the STRING PPI networks

We performed the network integration using an accuracy-based weighting method that uses the gene similarity scores for each gene interaction from the STRING PPI networks and the anatomy-based gene networks. This accuracy-based weighting method was used to integrate multiple networks in a previous work [50]. In this project, we used it to integrate only two network types: PPI and anatomy-based gene networks.

Initially, we evaluated the PPI and anatomy-based gene networks separately using the evaluation workflow described in the next section, which was used to decide the accuracy weights for the integration of gene similarity scores. For instance, if the accuracy values for the PPI and anatomy-based gene networks are *AC_1_* and *AC_2_* respectively, the weights for the PPI network (*W_1_*) and the anatomy-based gene network (*W_2_*) are calculated using equation 9 and equation 10, respectively. The accuracy values for each network are equivalent to their mean area under the curve (AUC) values of the receiver operating characteristic (ROC) curves generated during the evaluation.

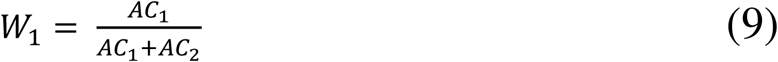

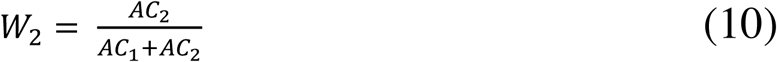

Then, the weights are used to calculate the gene similarity scores for the integrated network, based on the gene similarity scores of the original two networks. For instance, consider the similarity between the two genes: *G_a_* and *G_b_*. If the similarity scores from the PPI network and the anatomy-based gene network for those two genes are given by *sim1(G_a_, G_b_)* and *sim2(G_a_, G_b_)*, respectively, the similarity score in the integrated network: *sim3(G_a_, G_b_)* is calculated by the equation 11 below.

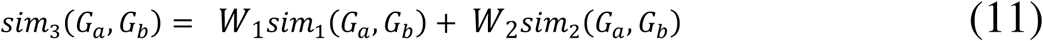

If an interaction is not found in an original network, the similarity score will be zero for that network; for instance, if *G_a_* and *G_b_* are not interacting in the PPI network, *sim1(G_a_, G_b_)* will be zero. During the calculation of weights, the most accurate original network will obtain a higher weight, and the integrated network will be weighted towards the more accurate network. We used this method to integrate the zebrafish and mouse PPI networks with the four anatomy-based gene networks (Lin, Resnik, Schlicker, and Wang), which resulted in four integrated networks for each organism.

### Evaluation of the network-based candidate gene prediction

For the evaluation, we used the reconciled zebrafish and mouse anatomical profiles initially obtained from the Monarch Initiative repository. We filtered the profiles to only keep the Uberon anatomy entities that contained at least 10 gene annotations. Then, we used leave-one-out cross-validation on one anatomical entity at a time and generated a ROC curve and a precision-recall curve for each term. Here, out of all the genes annotated to the anatomical entity of interest, the association of one gene is assumed to be unknown at one time, and the other genes are used to predict the anatomical term of that unknown gene. This process is repeated until all the genes are selected for anatomical term prediction. Then, we compared the distribution of area under the curve (AUC) values for all the anatomical entities for the three types of zebrafish and mouse networks (integrated networks, anatomy-based gene networks, and PPI networks) for each semantic similarity method (Lin, Resnik, Schlicker, and Wang). We used the Hishigaki method as the network-based candidate gene prediction method for which the equation 12 is given below.

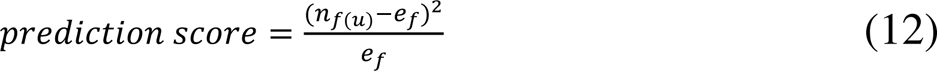

In the equation 12, *n_f(u)_* denotes the number of genes with the considered anatomical term (*f*) in the neighborhood of the gene in interest (*u*). Generally, the length of the neighborhood can be defined by the user but the immediate neighborhood (a length of one edge from the gene *u*) is shown to yield better results [51]; therefore, we only considered the immediate neighborhood of a gene for predictions. The expected frequency for the anatomical term is given by *e_f_*, which is calculated according to the equation 13 below.

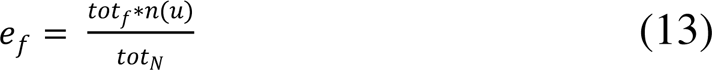

In the equation 13, *tot_f_* denotes the total number of genes annotated with the given anatomical term (*f*) in the network and *tot_N_* indicates the total number of genes in the network. The total number of genes in the immediate neighborhood of the gene of interest *(u)* is denoted by *n(u)*.

### Further validation of the prediction results

It is important to understand the biological significance of the evaluation results. For instance, during the previous step, if the integrated networks performed better than the PPI networks, it must be confirmed that the increased performance is due to the biological significance of integrating experimental anatomical data *via* anatomy ontology annotations and not due to random error/noise. For this purpose, we generated fully random networks with the same number of nodes and the same number of edges as those in the integrated and anatomy-based gene networks constructed using the Wang method. Furthermore, we generated another type of a random network by only randomizing the reconciled anatomical profiles by randomly assigning Uberon entities to each gene to match the original number of annotations, and then, constructing the anatomy-based gene networks and integrated networks using the Wang method. The second method is only a partial randomization because only the profiles were randomized, and the number of genes and the interactions are different from the original networks. From herein, we call the first random network type as ‘fully random networks’, and the second type as ‘random profile networks’. We compared the candidate gene prediction performances of the anatomy-based gene network and the integrated network for the Wang method with their corresponding fully random and random profile networks.

A potential concern with the proposed integrative method is that the same anatomical profiles used for the construction of the integrated networks were used for the evaluations. Therefore, if the integrated networks outperform the PPI network, this may have been caused by the circular use of the same anatomical profile for the network construction and the evaluation. To further investigate this issue, we conducted two experiments. First, we randomly removed 30 Uberon entities with at least 10 gene annotations from the reconciled zebrafish anatomical profiles and constructed the anatomy-based and integrated networks from the remaining anatomical entities using the Wang method. Then we compared the network-based candidate gene prediction performance of those networks with the zebrafish PPI network using the removed 30 Uberon entities for the evaluation. If the performance increase in the integrated network is only due to re-using the same Uberon entities for the evaluation, the integrated network should not show a performance increase compared to the PPI network when those 30 Uberon entities were used for the evaluation because they were not involved in the network construction.

For the second experiment, we downloaded zebrafish Gene Ontology (GO) annotations from GO consortium and pre-processed them to keep only the Gene Ontology-Biological Process (GO-BP) annotations. Then, we constructed GO-BP profiles for zebrafish genes and reconciled them to only keep the genes that were found in zebrafish PPI, anatomy-based gene networks, and integrated networks. Finally, we evaluated the network-based candidate gene prediction performance of the anatomy-based and integrated networks constructed using the Wang method and the zebrafish PPI network using the reconciled GO-BP profiles. Here, evaluation was performed by GO-BP profiles, which were not used for the construction of the anatomy-based and integrated networks.

## Results

### Data Sources

The raw zebrafish STRING PPI network contained 23,018 genes and 12,558,675 interactions and the mouse STRING PPI network contained 21,052 genes and 6,262,253 interactions. After applying the 0.7 gene similarity score cutoff to keep only the high-quality interactions, the filtered zebrafish PPI network contained 14,677 genes and 501,704 interactions, and the filtered mouse PPI network contained 13,866 genes and 414,667 interactions.

The original zebrafish anatomical profiles retrieved from the Monarch Initiative repository contained 5,405 genes annotated to 960 Uberon anatomy entities, and the mouse profiles contained 14,652 genes annotated to 1,537 Uberon entities (Table 1). Not all of these genes were found in the STRING PPI networks, owing to differences in the data sources. After implementing the gene reconciliation algorithm that contained three rounds (direct name matching, Ensembl ID matching, and gene synonym matching), the number of original matches was increased from 2,527 (direct name matching) to 3,048 for the zebrafish and from 8,166 (direct name matching) to 8,607 for the mouse (Table 1). The detailed reconciliation statistics are shown in Table 1.

**Table 1.** The statistics for the reconciliation of gene names between the anatomical profiles from Monarch, and the PPI networks from STRING, for zebrafish and mouse.

|  | Zebrafish | Mouse |
| --- | --- | --- |
| Number of genes in the original anatomical profiles | 5,405 | 14,652 |
| Number of Uberon entities in the original anatomical profiles | 960 | 1,537 |
| Number of genes directly matched to the PPI network using the gene name (round 1) | 2,527 | 8,166 |
| Number of genes matched using the ensemble IDs (round 2) | 402 | 378 |
| Number of genes matched using synonyms in the STRING database (round 3) | 119 | 63 |
| Number of final gene matches to the PPI network (in all 3 rounds)/ Number of genes in the reconciled anatomical profile | 3,048 | 8,607 |
| Number of Uberon entities in the reconciled anatomical profile | 943 | 1,524 |
| Number of final gene mismatches | 2,357 | 6,045 |
| Number of genes kept in the anatomical profiles after filtering out the Uberon entities with fewer than 10 gene annotations | 3,037 | 8,606 |
| Number of Uberon entities kept in the anatomical profiles after filtering out the Uberon entities with fewer than 10 gene annotations | 294 | 850 |

The extra 521 genes for the zebrafish and 441 genes for the mouse that were matched during round 2 (using Ensembl ids) and round 3 (using gene synonyms) contained outdated gene names in the PPI networks. Therefore, they were updated to the correct names that were used in the anatomical profiles. The final number of gene mismatches for zebrafish and mouse are 2,357 and 6,045, respectively. The majority of these mismatched genes have anatomical term annotations in the Monarch Initiative repository, but do not have a characterized protein. Therefore, there are no protein interaction records in the STRING database. After the reconciliation, the original anatomical profiles were filtered to contain only the genes that were matched with the PPI networks and to contain only the Uberon entities that have at least 10 gene annotations (Table 1). These reconciled and filtered anatomical profiles were used during the evaluation of different network types because it is important to evaluate the networks using the genes that are found in all the three types of networks (PPI, anatomy-based gene networks, and integrated networks) for zebrafish and mouse for a valid comparison.

### Construction of the anatomy-based gene networks

When constructing the anatomy-based gene networks, we used original anatomical profiles (before the reconciliation) to retain all of their genes in the networks. We used the reconciled anatomical profiles only for the evaluation of the networks. The gene similarity score distributions for the four types of unfiltered anatomy-based gene networks (Lin, Resnik, Schlicker, and Wang) for the zebrafish and the mouse are shown in Figure 3 and Figure 4, respectively. The gene similarity scores for Lin, Resnik, and Schlicker methods are distributed approximately between a range from 0 to 0.40. In contrast, the distribution for the Wang method is symmetrical around the 0.50 region.

**Fig. 3.**
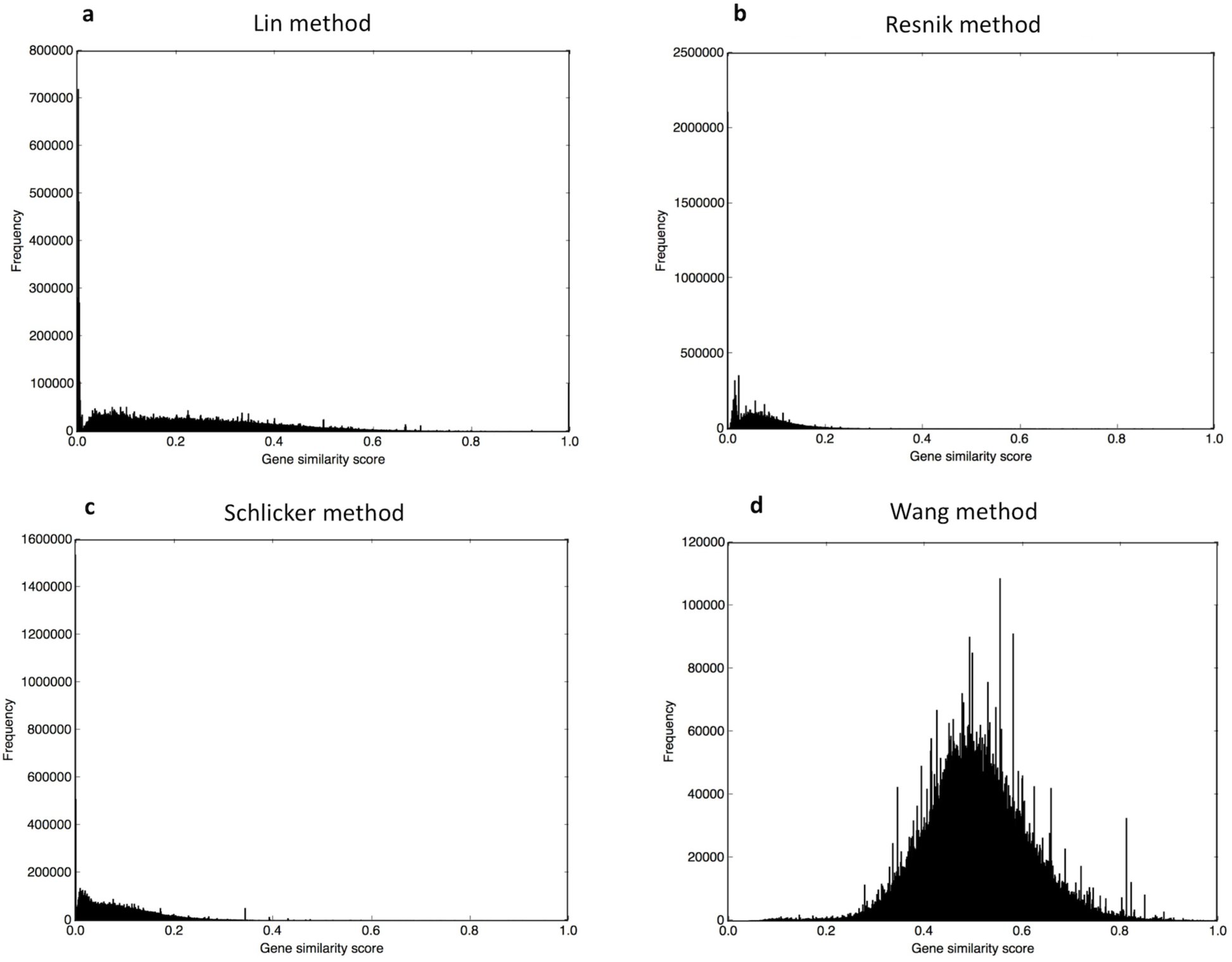
The gene similarity score distributions for the zebrafish unfiltered anatomy-based gene networks constructed by (a) Lin method, (b) Resnik method, (c) Schlicker method, and (d) Wang method.

**Fig 4.**
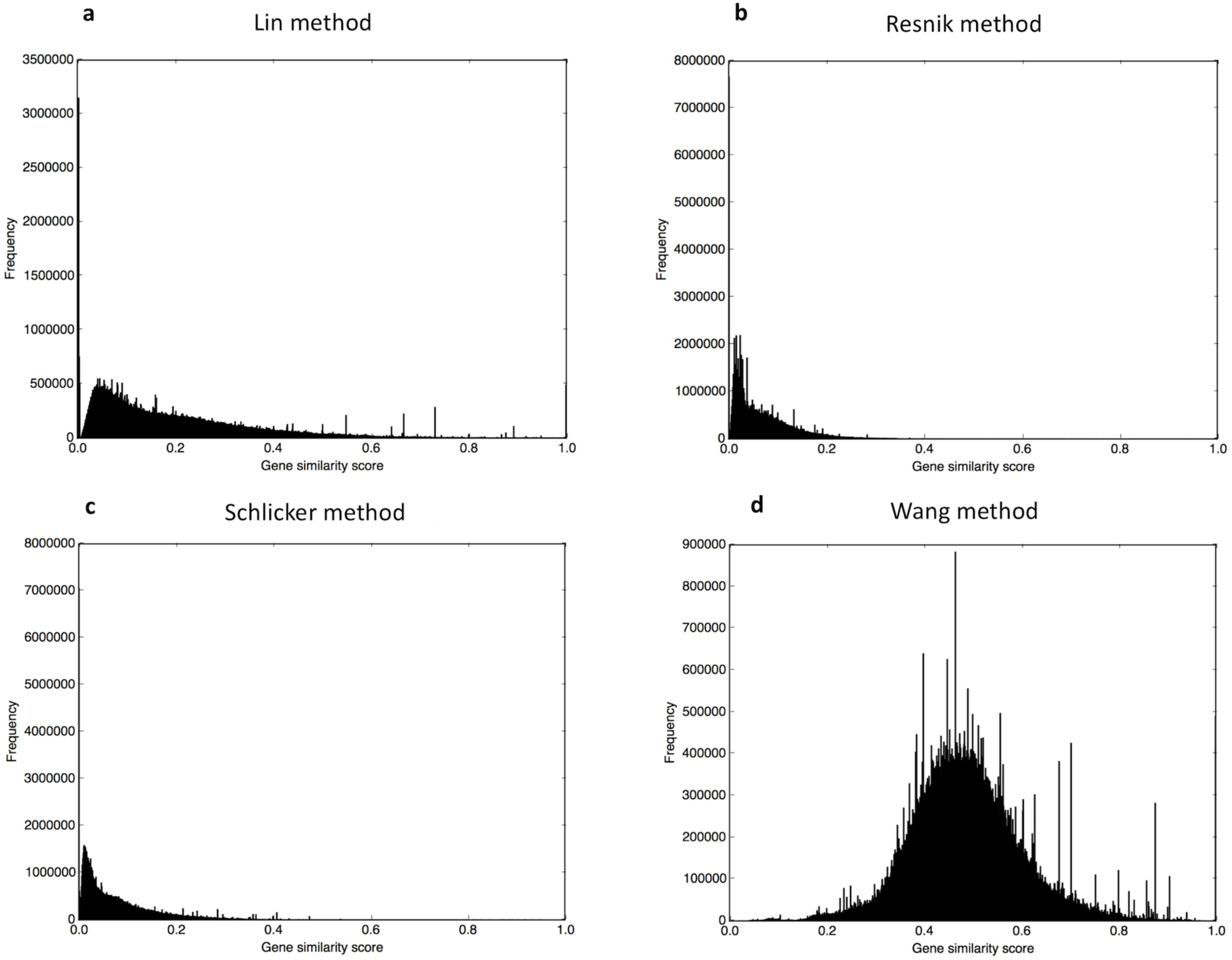
The gene similarity score distributions for the mouse unfiltered anatomy-based gene networks constructed by (a) Lin method, (b) Resnik method, (c) Schlicker method, and (d) Wang method.

Obtaining these distributions was critical to determine the gene similarity score cutoff applied to each network. For example, applying 0.7 as the cutoff for the Wang anatomy-based gene network for the zebrafish, generated a filtered network with 5,386 genes and 789,282 interactions; if the same 0.7 cutoff was applied to the zebrafish Resnik network, the filtered network would only have 30 genes and 31 interactions. If these two networks were evaluated, the changes in the number of genes and the number of interactions would have a significant effect on their performance. Therefore, a cutoff must be applied to keep the network size relatively constant among the different networks. However, it is difficult in practice to apply cutoffs to keep the exact number of genes and the interactions among the networks. Therefore, using the trial and error method, we applied different cutoffs to anatomy-based gene networks to keep the number of interactions between 500,000 and 750,000. The statistics for the network sizes of filtered and unfiltered networks and their cutoffs are shown in Table 2 and Table 3 for the zebrafish and the mouse, respectively.

**Table 2.** The statistics for the unfiltered and filtered anatomy-based gene networks for zebrafish

|  | Lin method | Resnik method | Schlicker method | Wang method |
| --- | --- | --- | --- | --- |
| Number of genes in the unfiltered network | 5,405 | 5,405 | 5,405 | 5,405 |
| Number of interactions in the unfiltered network | 14,534,897 | 14,534,897 | 14,534,897 | 14,604,258 |
| The gene similarity score cutoff | 0.55 | 0.18 | 0.24 | 0.70 |
| Number of genes in the filtered network | 5,387 | 4,909 | 5,401 | 5,386 |
| Number of interactions in the filtered network | 700,138 | 712,286 | 692,539 | 789,282 |

**Table 3.** The statistics for the unfiltered and filtered anatomy-based gene networks for mouse

|  | Lin method | Resnik method | Schlicker method | Wang method |
| --- | --- | --- | --- | --- |
| Number of genes in the unfiltered network | 14,652 | 14,652 | 14,652 | 14,652 |
| Number of interactions in the unfiltered network | 107,094,117 | 107,094,117 | 107,094,117 | 107,324,905 |
| The gene similarity score cutoff | 0.9 | 0.32 | 0.41 | 0.95 |
| Number of genes in the filtered network | 9,784 | 10,081 | 12,755 | 9,126 |
| Number of interactions in the filtered network | 588,359 | 536,602 | 522,183 | 510,139 |

### Integration of the anatomy-based gene networks with the STRING PPI networks

During the integration, we combined unfiltered anatomy-based gene networks for zebrafish and mouse with the corresponding STRING PPI networks. When selecting the gene similarity cutoffs for filtering the integrated networks, we considered their gene similarity score distributions as explained in the previous section. The statistics for the filtered and unfiltered network sizes are shown in Table 4 for the zebrafish and Table 5 for the mouse. The generated integrated networks are larger than the anatomy-based gene networks in terms of the number of genes and the interactions. For instance, the Wang anatomy-based gene network for the zebrafish has 5,405 genes and 14,604,258 interactions (Table 2), whereas the zebrafish integrated network for the Wang method has 25,375 genes and 26,821,274 interactions (Table 4). During the integration, the 5,405 genes in the anatomy-based gene network were integrated with the 23,018 genes in the zebrafish PPI network, which caused an increase in the network size. The common genes and interactions were retained according to the integration formula (equation 11), and the genes and the interactions that are unique to one network were also included in the integrated network if the final gene similarity score is above the cutoff. Therefore, the integrated networks are more complete in terms of the number of genes and the information contained.

**Table 4.** The statistics for the unfiltered and filtered integrated networks for zebrafish

|  | Lin method | Resnik method | Schlicker method | Wang method |
| --- | --- | --- | --- | --- |
| Number of genes in the unfiltered network | 25,375 | 25,375 | 25,375 | 25,375 |
| Number of interactions in the unfiltered network | 26,753,086 | 26,753,086 | 26,753,086 | 26,821,274 |
| The gene similarity score cutoff | 0.33 | 0.23 | 0.24 | 0.4 |
| Number of genes in the filtered network | 17,394 | 20,066 | 20,929 | 13,940 |
| Number of interactions in the filtered network | 730,855 | 726,589 | 690,208 | 744,519 |

**Table 5.** The statistics for the unfiltered and filtered integrated networks for mouse

|  | Lin method | Resnik method | Schlicker method | Wang method |
| --- | --- | --- | --- | --- |
| Number of genes in the unfiltered network | 27,097 | 27,097 | 27,097 | 27,097 |
| Number of interactions in the unfiltered network | 111,461,010 | 111,461,010 | 111,461,010 | 111,690,355 |
| The gene similarity score cutoff | 0.50 | 0.27 | 0.30 | 0.53 |
| Number of genes in the filtered network | 13,125 | 17,898 | 18,002 | 12,916 |
| Number of interactions in the filtered network | 653,848 | 661,619 | 613,671 | 712,720 |

The gene similarity score distributions for the integrated networks for zebrafish and mouse are shown in Figure 5 and Figure 6, respectively. When compared to the distributions of the corresponding anatomy-based gene networks as shown in Figures 3 and 4, the distributions of the integrated networks are slightly skewed to the right; especially, the gene similarity scores of the Wang anatomy-based gene networks are symmetrical and distributed around 0.5; in contrast, the distributions for the Wang integrated networks are shifted to 0-0.50 region. This is due to the effect of the integration. Only the interactions that have high similarity scores in the anatomy-based gene network and the PPI network receive higher scores in the integrated network. Most of the interactions in the anatomy-based gene network received low support from the PPI network, thus the gene similarity score distribution of the integrated network is skewed to right. By applying the cutoffs as shown in Tables 4 and 5, the interactions with the highest similarity scores, which received support from both the PPI and the anatomy-based gene networks, could be selected.

**Fig. 5.**
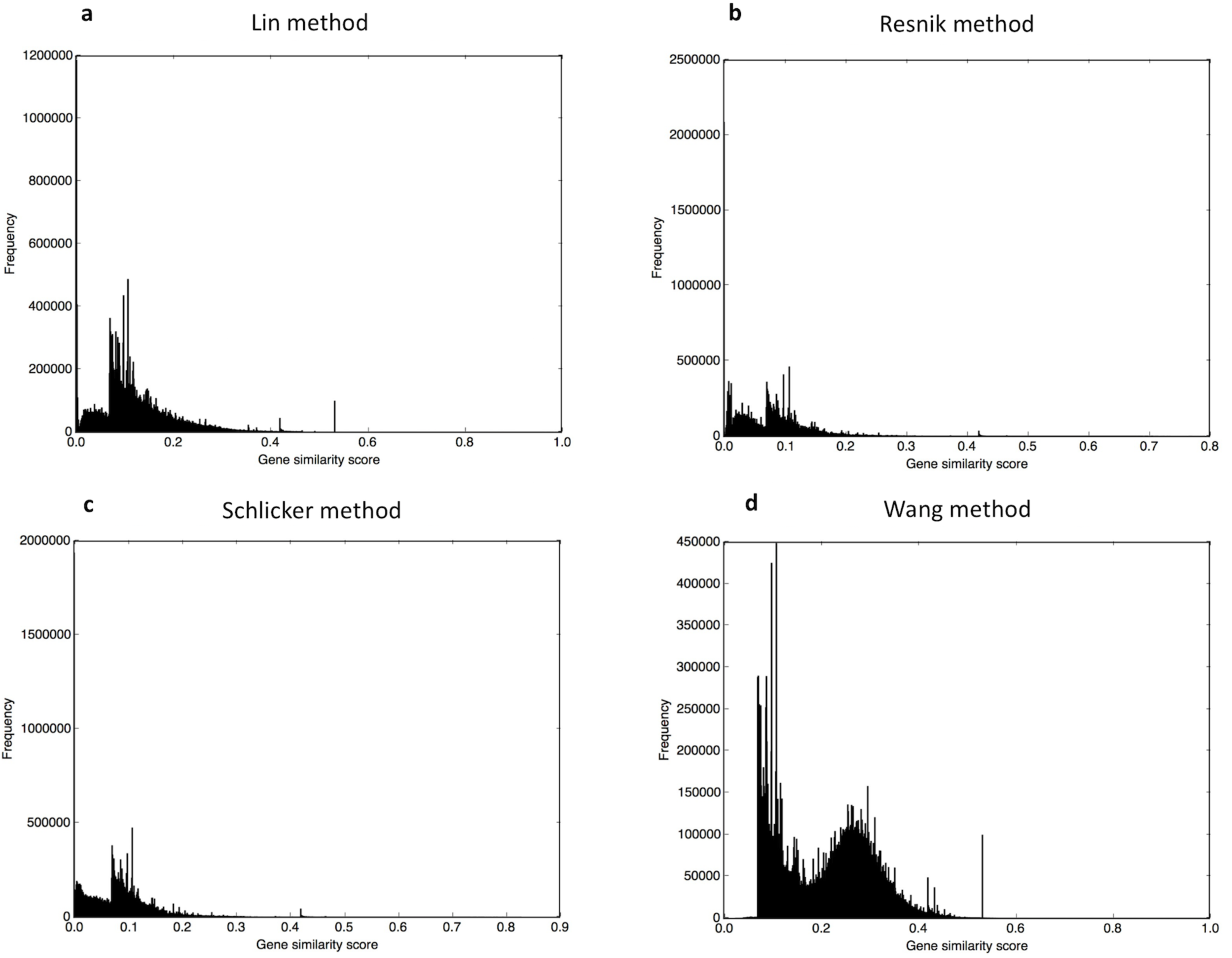
The gene similarity score distributions for the zebrafish unfiltered integrated networks constructed by (a) Lin method, (b) Resnik method, (c) Schlicker method, and (d) Wang method.

**Fig. 6.**
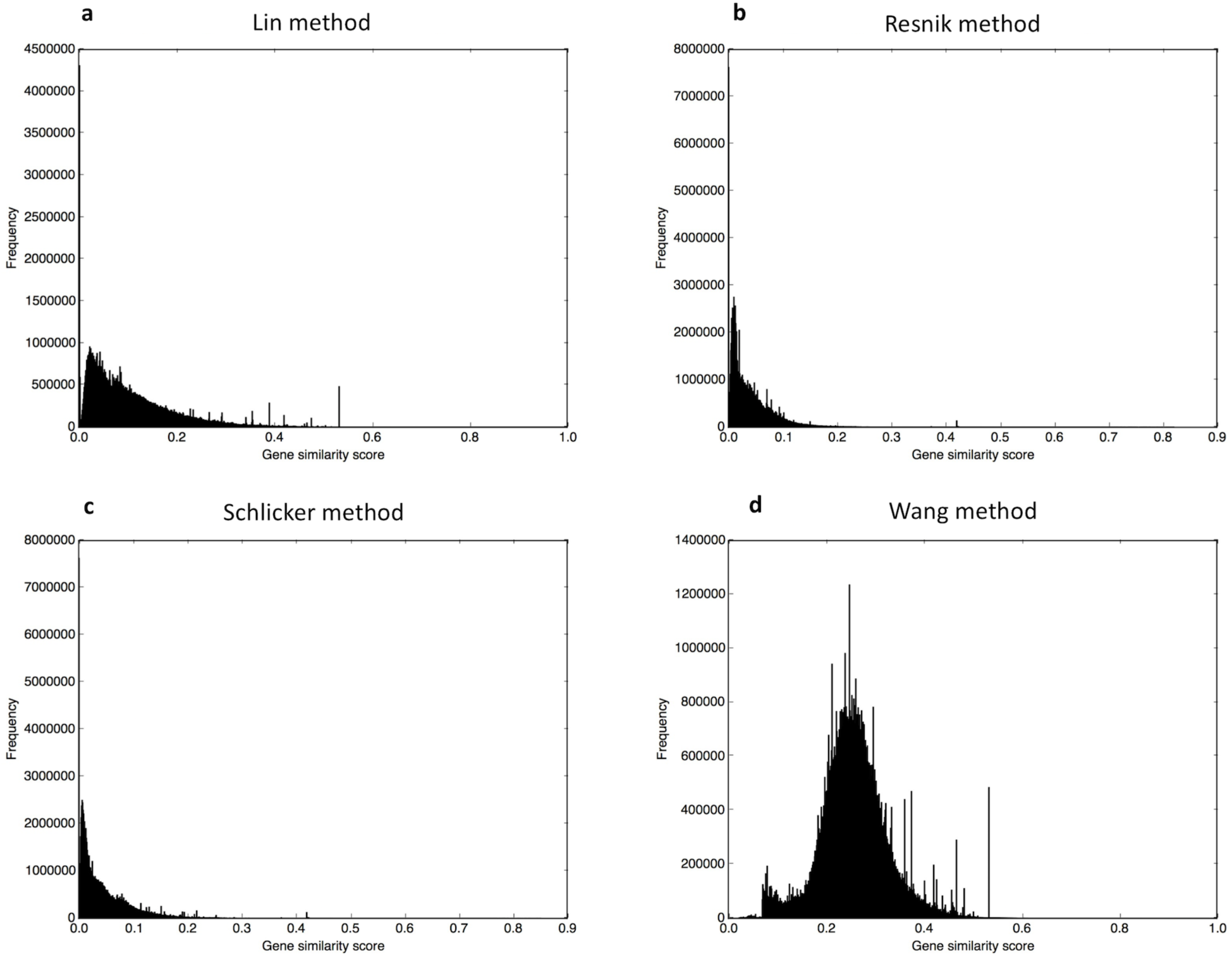
The gene similarity score distributions for the mouse unfiltered integrated networks constructed by (a) Lin method, (b) Resnik method, (c) Schlicker method, and (d) Wang method.

### Evaluation of the candidate gene predictions

The purpose of integrating anatomy ontology data with PPI networks was to improve the accuracy of predicting candidate genes for anatomical entities. The boxplot comparisons of the AUC distributions of ROC and precision-recall curves for zebrafish networks are given in Figure 7 and Figure 8, respectively. The boxplot comparisons of the AUC distributions of ROC and precision-recall curves for mouse networks are given in Figure 9 and Figure 10, respectively. Each figure compares AUC distributions for three network types: anatomy-based gene networks, integrated networks, and PPI networks for the four semantic similarity calculation methods.

**Fig. 7.**
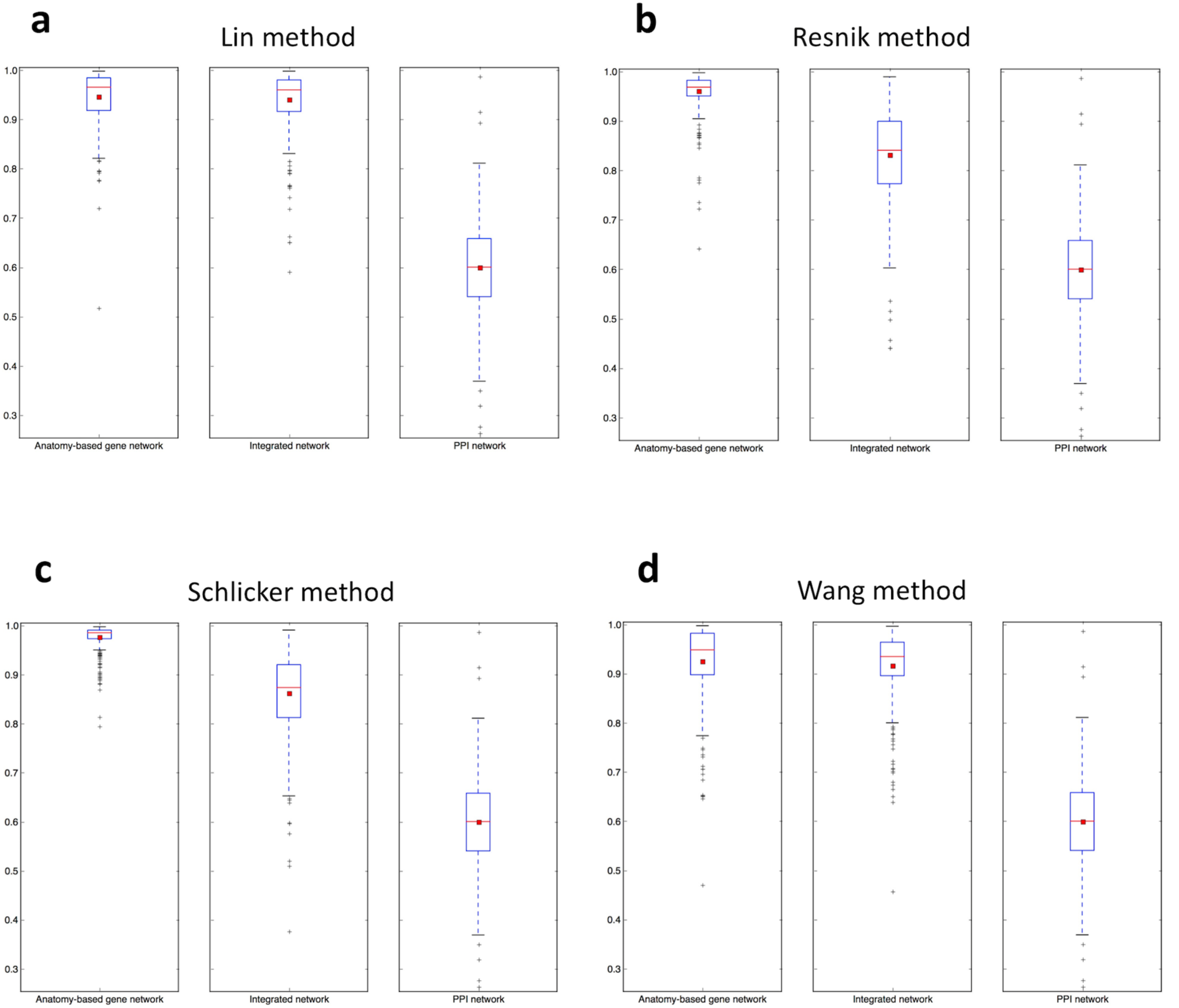
The boxplot comparisons for the AUC distributions of ROC curves for filtered anatomy-based gene networks, integrated networks, and PPI networks for the four semantic similarity calculation methods for the zebrafish. In the boxplots, the red line and the square represent the median and mean, respectively.

**Fig. 8.**
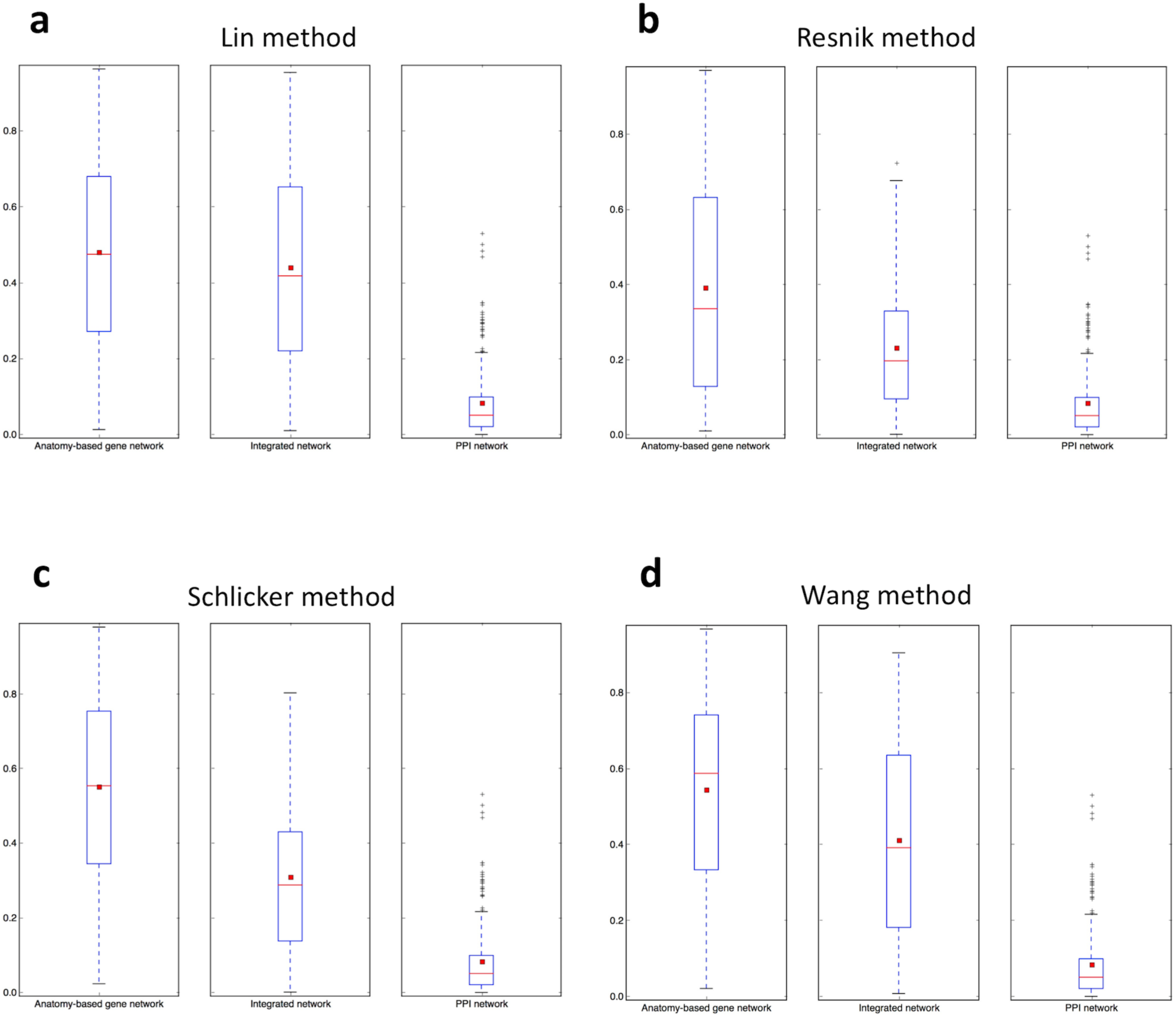
The boxplot comparisons for the AUC distributions of precision-recall curves for filtered anatomy-based gene networks, integrated networks, and PPI networks for the four semantic similarity calculation methods for the zebrafish. In the boxplots, the red line and the square represent the median and mean, respectively.

**Fig. 9.**
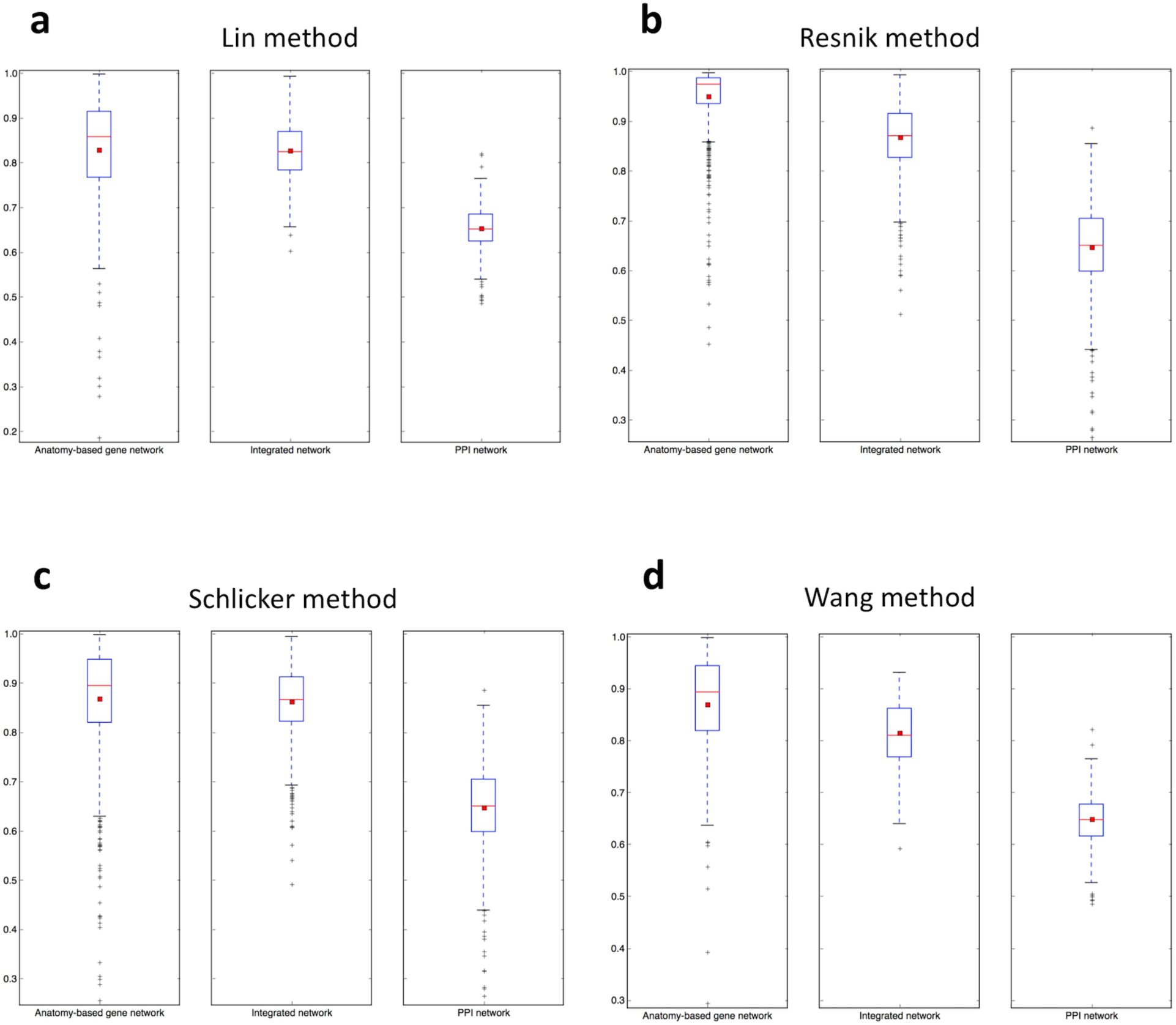
The boxplot comparisons for the AUC distributions of ROC curves for filtered anatomy-based gene networks, integrated networks, and PPI networks for the four semantic similarity calculation methods for the mouse. In the boxplots, the red line and the square represent the median and mean, respectively.

**Fig. 10.**
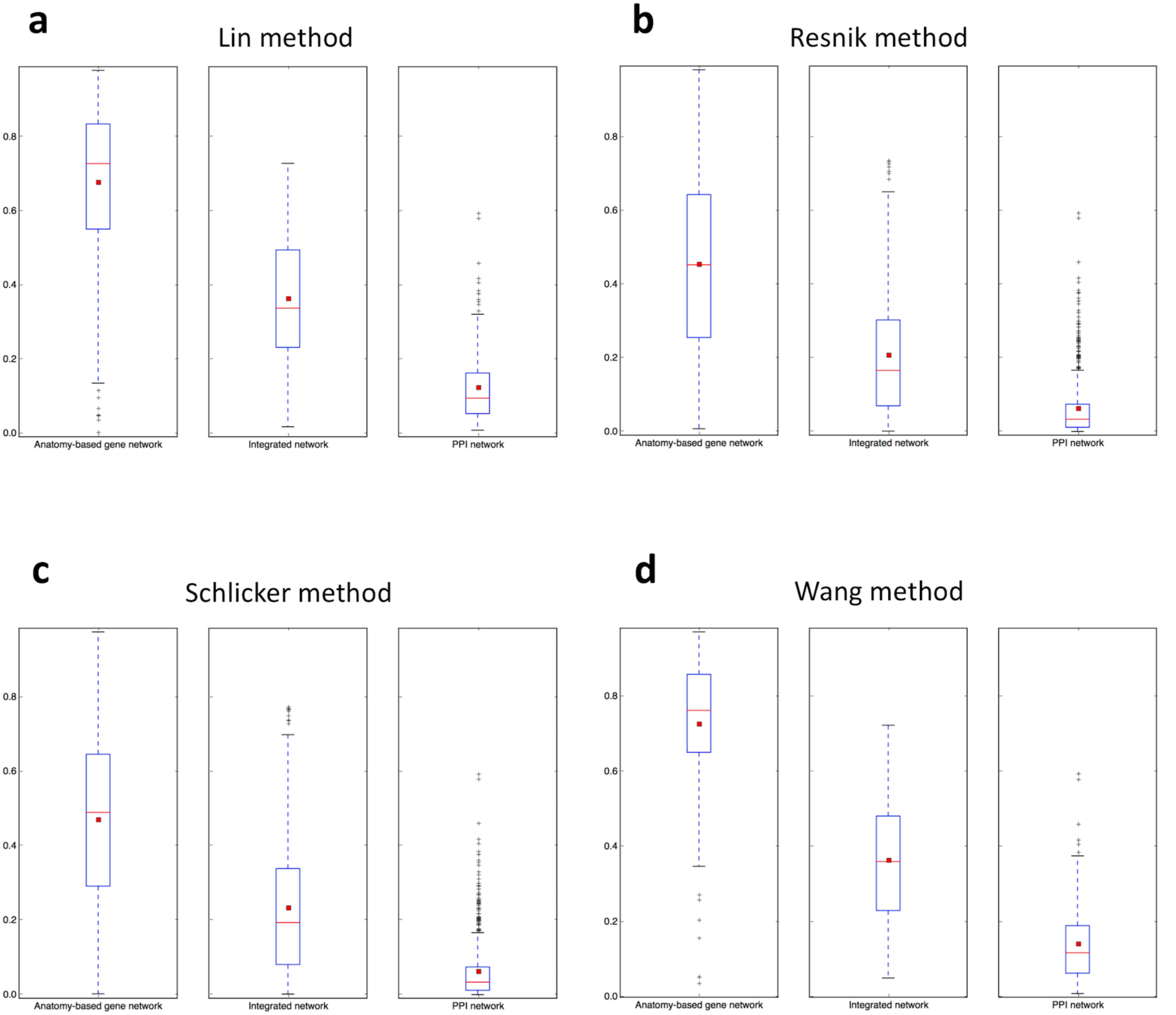
The boxplot comparisons for the AUC distributions of precision-recall curves for filtered anatomy-based gene networks, integrated networks, and PPI networks for the four semantic similarity calculation methods for the mouse. In the boxplots, the red line and the square represent the median and mean, respectively.

According to the boxplot comparisons, the AUC distributions of ROC and precision-recall curves for the integrated networks are higher than the PPI networks for both zebrafish and mouse for all four semantic similarity calculation methods. This confirms that integration of anatomy ontology data improves the candidate gene prediction accuracy for anatomical entities compared to the PPI networks for two model organisms. Although anatomy-based gene networks outperform integrated networks in the boxplot comparisons, integrated networks are more suitable for candidate gene prediction for anatomical entities in practice. For instance, anatomy-based gene networks are much smaller (incomplete) than integrated networks, thus the genes that can be predicted using anatomy-based gene networks are very limited. Moreover, integrated networks include support from multiple data sources such as anatomical annotations and other molecular interactions, thus improving the prediction power.

### Further validation of the prediction results

The AUC distribution comparisons of ROC and precision-recall curves for the non-randomized anatomy-based gene network and the integrated network for the Wang method with their fully random and random profile counterparts are shown in Figure 11.

**Fig. 11.**
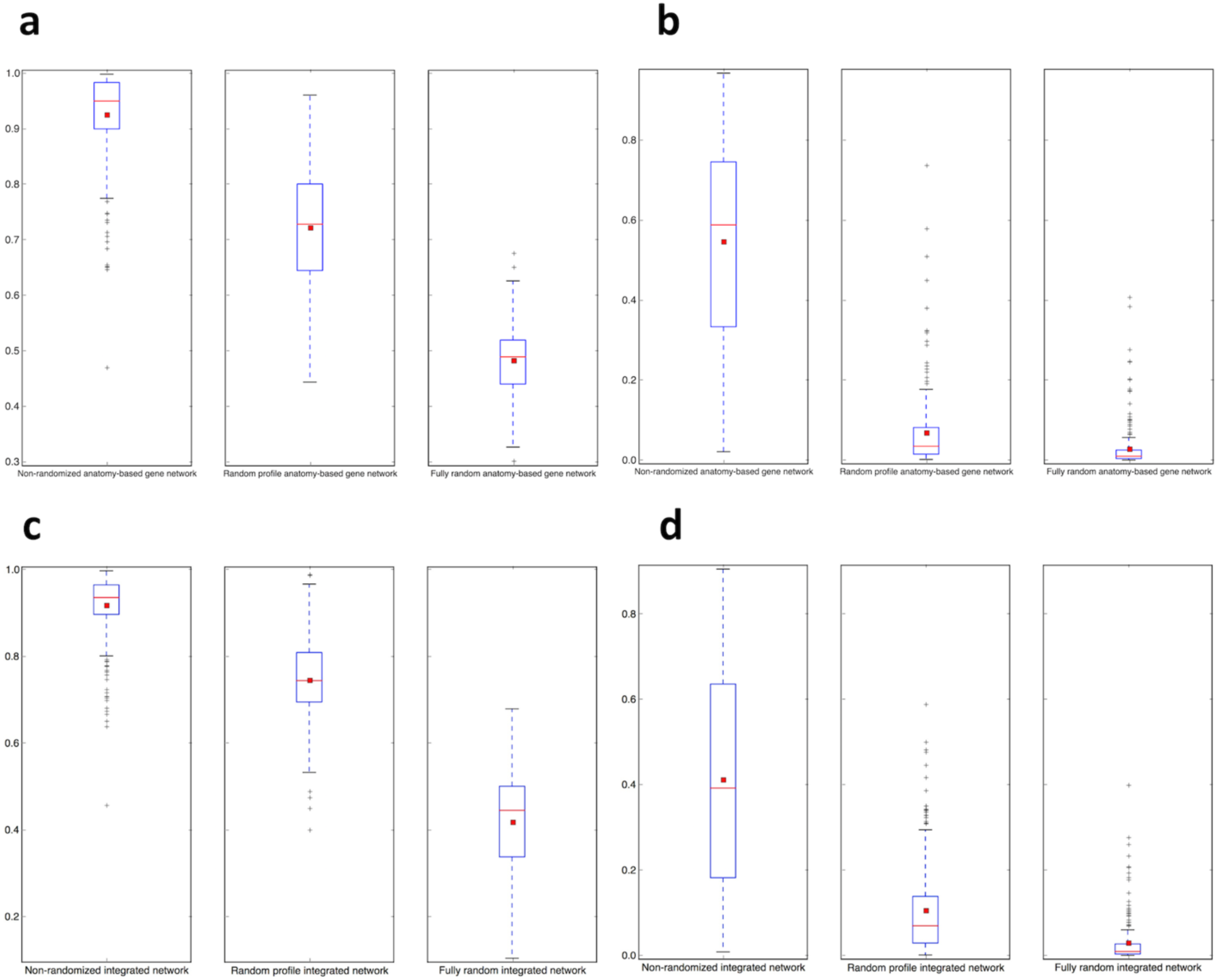
The boxplot comparisons of the AUC distributions for (a) ROC and (b) precision-recall curves for the filtered non-randomized anatomy-based gene network, random profile anatomy-based gene network, and fully random anatomy-based gene network for the Wang method for the zebrafish. The boxplot comparisons of the AUC distributions for (c) ROC and (d) precision-recall curves for the filtered non-randomized integrated network, random profile integrated network, and fully random integrated network for the Wang method for the zebrafish. In the boxplots, the red line and the square represent the median and mean, respectively.

According to the comparisons, non-randomized networks have a higher performance (AUC score distributions) compared to the randomized networks in both network types. When comparing the two randomization methods, random profile networks, which were constructed by only randomizing the anatomy profiles have a higher performance than the fully random networks. This comparison with randomized networks indicates that the high performance observed by the anatomy-based gene network and the integrated network is due to their biological significance.

To test the effect of the circular use of the same anatomy profiles for anatomy-based gene network construction and for the network evaluation, the AUC distribution comparisons of ROC and precision-recall curves for Wang anatomy-based and integrated networks evaluated using 30 Uberon anatomy entities that were not used for the construction of the networks are given in Figure 12.

**Fig. 12.**
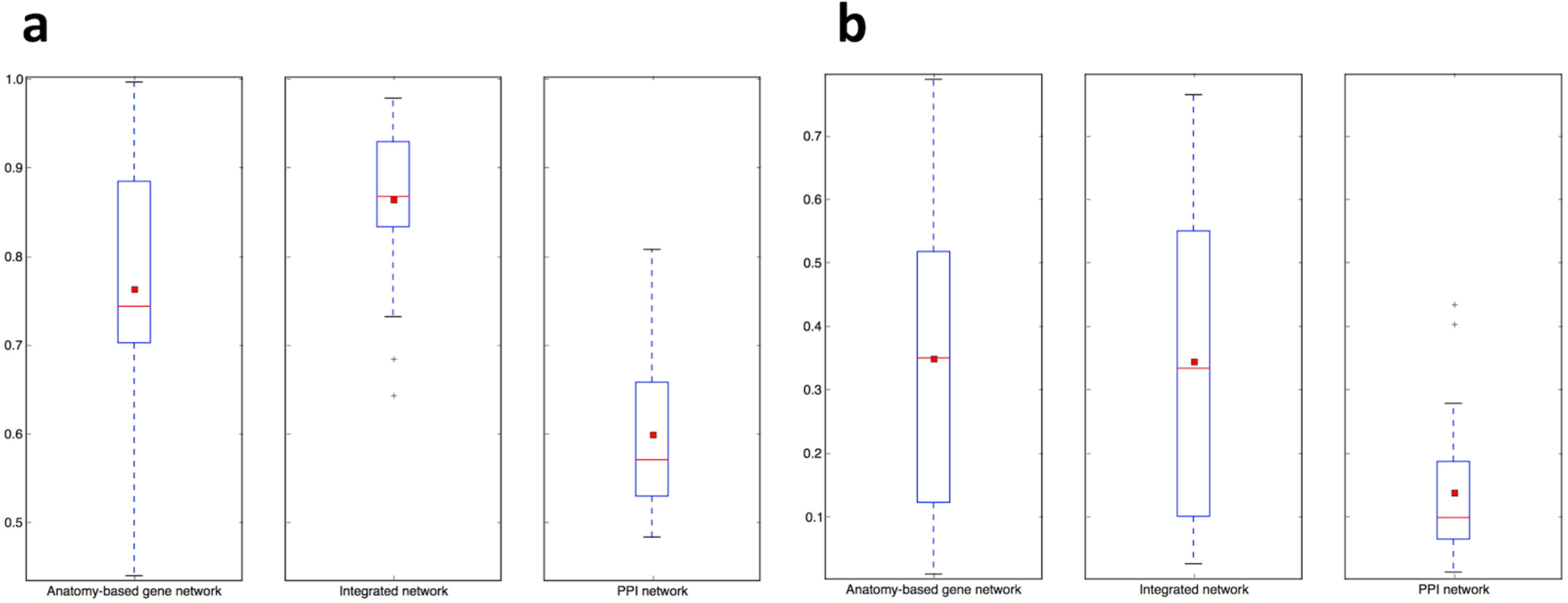
The boxplot comparisons of the AUC distributions for (a) ROC and (b) precision-recall curves for the filtered integrated network, PPI network, and anatomy-based gene network using the Wang method for the zebrafish. The integrated network and the anatomy-based gene network were generated using the zebrafish anatomy profile after randomly removing 30 Uberon entities, which had at least 10 gene annotations. The same 30 entities were used for the evaluation. In the boxplots, the red line and the square represent the median and mean, respectively.

According to Figure 12, particularly based on AUC distribution comparison for ROC curves, the integrated network performs better than the PPI network, even when evaluated using the 30 Uberon entities that were not used for the network construction. Here, the integrated network has a better performance than the anatomy-based gene network.

The AUC distribution comparisons of ROC and precision-recall curves for Wang anatomy-based and integrated networks, evaluated using GO-BP profiles for the zebrafish are shown in Figure 13. Because the anatomy-based gene network and the integrated network were constructed using the zebrafish anatomy profile using the Wang method, the GO-BP annotations, which were used for the evaluation, do not have a direct influence on the network construction. The integrated network performs better than both the PPI and anatomy-based gene networks, when evaluated by the GO-BP profiles (Figure 13).

**Fig. 13.**
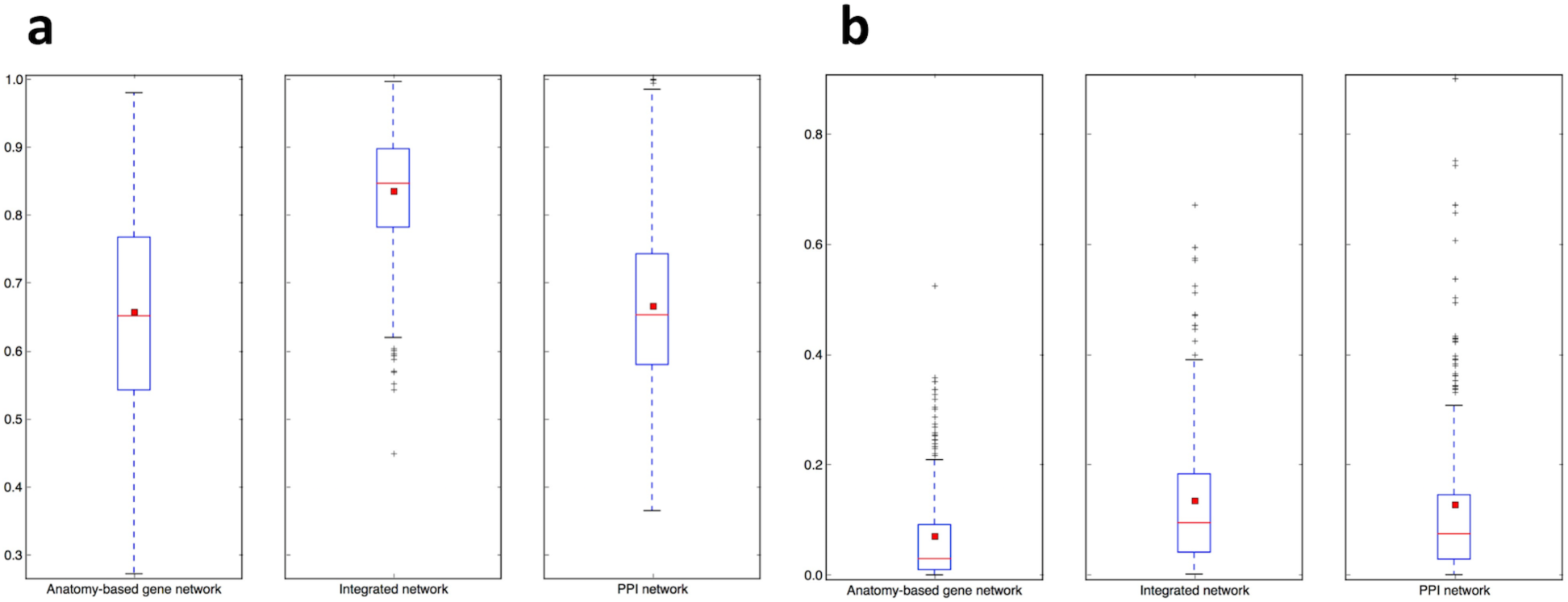
The boxplot comparisons of the AUC distributions of (a) ROC and (b) precision-recall for the filtered integrated network, PPI network, and anatomy-based gene network for the Wang method in zebrafish. The networks were evaluated using the annotation profiles containing Gene Ontology-Biological Process (GO-BP) entities for the zebrafish genes. In the boxplots, the red line and the square represent the median and mean, respectively.

## Discussion

The goal of this work was to test whether the integration of anatomy ontology data with PPI networks enhances the network-based candidate gene prediction accuracy when predicting gene candidates associated with anatomical entities. Discovering new genes associated with anatomical phenotypes, such as limb and fin development leads to a clear understanding of the underlying molecular mechanisms for anatomical development. Moreover, such discoveries support the progress of emerging evolutionary developmental biology (evo-devo) studies [2, 52]. Computational prediction of candidate genes for anatomical entities is rare compared to predicting genes for molecular mechanisms and functions, but the need for accurate methods is evident based on the low number of gene annotations currently available for the majority of anatomical entities. This led us to utilize existing network-based candidate gene prediction methods and improve the candidate gene prediction accuracy by integrating anatomy ontology data with the existing PPI networks.

The candidate gene predictions based on PPI networks suffer from low prediction accuracy due to the low quality of the networks [13, 18–21]. One potential way to increase their quality is to incorporate the experimental knowledge about gene-anatomical entity relationships that is extracted from the literature into PPI networks. Here, using the Uberon anatomy ontology and semantic similarity calculation methods, we developed a computational framework to integrate experimental knowledge about gene-anatomical entity associations with PPI interaction data. First, we constructed anatomy-based gene networks by calculating pairwise semantic similarity scores between genes using their respective anatomical profiles, which represents a network of genes based on their Uberon annotations. In this network, if two genes receive a high gene similarity score, they are associated with similar anatomical entities. Next, we integrated the anatomy-based gene networks for the mouse and the zebrafish with the corresponding PPI networks downloaded from the STRING database. In the integrated networks, the gene interactions that receive high interaction scores are the ones that receive a higher support from both the PPI and anatomy-based gene networks. This filters out false positive interactions in the PPI networks if those interactions are not supported by the anatomy-based gene networks, that is, if they receive low scores from the anatomy-based gene networks. On the other hand, gene interactions with low scores in the PPI networks will be enhanced if the interactions are supported by the anatomy-based gene networks, that is, if they have high similarity scores from the anatomy-based gene networks. In cases where the gene similarity score is zero in anatomy-based gene networks due to lack of anatomical term annotations, the support from the PPI network should be extremely high for those interactions to receive a moderate similarity score in the integrated networks and potentially be selected after application of the gene similarity score cutoff. Similarly, if two proteins are not interacting in a PPI network, they need a higher support from the anatomy-based gene network to receive a moderate gene similarity score in the integrated network and to be selected after the application of the gene similarity score cutoff. Furthermore, a protein usually will interact with multiple proteins, thus has a degree larger than two in the graph. The integrated networks not only increase the network size, but also increase the average degrees of proteins in the graph, which improves the network completeness.

According to the network performance evaluations (Figures 7, 8, 9, and 10), the integrated networks perform better than the PPI networks, showing that the integration increases their quality. The robustness of this conclusion was further demonstrated by performing evaluations under different settings. For example, we used both ROC curves and precision-recall curves for the evaluations. Moreover, we used four semantic similarity methods (Lin, Resnik, Schlicker, and Wang) to construct semantic networks for the two model organisms (zebrafish and mouse). Under all these experimental settings, the integrated networks performed better than the PPI networks, strengthening the conclusion that the integration of the literature knowledge that has been annotated using an anatomy ontology with the PPI networks increases the candidate gene prediction accuracy for anatomical entities.

To test the biological significance of the results, we compared the integrated and the anatomy-based gene networks constructed using the Wang method for the zebrafish with their randomized counterparts (Figure 11). The results demonstrated that the higher candidate gene prediction performance observed in the integrated network has biological significance and is not due to random error/chance. Of the two randomization procedures used, the random profile networks that were generated by only randomizing the anatomical profiles, perform better than the fully random networks that were constructed by completely randomizing the entire networks. When only the anatomical profiles are randomized, the original number of annotations per gene was kept constant even after the randomization, which may lower the randomization effect by including closely related Uberon entities for the same gene. This may explain their higher performance compared to the fully random networks. However, the non-randomized networks have a higher performance than the two randomized network types, especially, in precision-recall curves, due to their biological significance.

Another challenge we faced during the analysis is the concern of circular use of the same anatomical profiles for the construction of the networks (anatomy-based gene networks and then the integrated networks) and for their evaluation. The increased performance observed in the integrated networks may be due to using the same anatomical profiles for the evaluation. We conducted two experiments to assess whether the circular use of the anatomical profiles affects the observed results. First, we randomly selected 30 Uberon entities and removed their annotations from the zebrafish anatomical profiles and constructed the networks using the Wang method, which were evaluated using the same 30 entities (Figure 12). Second, we evaluated the networks using the GO-BP profiles for the zebrafish (Figure 13). In both experiments, the annotations used for the network construction were not used for the evaluation, and in both the occasions the integrated network outperformed both the PPI and anatomy-based gene networks. This indicates the increased performance observed in the integrated networks is not due to the circular use of the same anatomical profiles.

Previous studies [17, 36, 37] that used semantic networks for candidate gene predictions tend to use networks that are directly based on the ontology for candidate gene predictions. However, the works Jiang, et al. [37] and Cho, et al. [38] suggest the integration of semantic networks with PPI networks as a possible method to improve the candidate gene prediction accuracy. Therefore, we integrated the anatomy-based gene networks that were constructed using anatomy ontology annotations with PPI networks. Using the integrated networks over the anatomy-based gene networks have several advantages. First, anatomy-based gene networks contain a limited number of genes that only have Uberon annotations, thus they are not as practical for candidate gene prediction as the integrated networks. For example, the zebrafish filtered anatomy-based gene network constructed using the Schlicker method contains 5,401 genes (Table 2), whereas the corresponding integrated network contains 20,929 genes (Table 4). Integration of PPI network data adds new genes without anatomical annotations to the integrated networks; these genes are then novel potential candidates for anatomical entities. Furthermore, the anatomy-based gene networks represent the gene organization in the network based only on their anatomical annotations, and they do not include the molecular interactions coming from experimental sources, which is achieved after the integration.

We sought to understand our initial finding that the anatomy-based gene networks outperformed both integrated and PPI networks during the initial evaluations (Figures 7, 8, 9, and 10). For these evaluations, we used the same anatomical profiles for both the construction and the evaluation of the anatomy-based and integrated networks. This was essential to capture the maximum anatomical information available in the anatomical profiles during the network construction. However, in a practical scenario, these semantic networks are used to predict novel gene-anatomical entity associations, which are not available for the construction of these networks. This scenario is simulated when we removed annotations for 30 Uberon entities from zebrafish anatomical profiles and used the remaining annotations for the construction of semantic networks; subsequently, used the removed annotations for the evaluations (Figure 12). Here, the integrated network outperforms both the anatomy-based gene network and PPI network. Furthermore, when we used GO-BP profiles to evaluate the networks, again, the integrated network outperforms the other two networks. This indicates, when the circular use of the same anatomical profiles for the construction and the evaluation of the semantic networks is eliminated, the integrated network has the best performance. Because of aforementioned reasons, integrated networks are better suited for prediction of new gene candidates for anatomical entities than the anatomy-based gene networks.

### Conclusions and Future work

This work focuses on improving the quality of the PPI networks by integrating known gene-anatomy term knowledge *via* anatomy ontology data. According to candidate gene prediction performance evaluations tested under different computational settings (four semantic similarity calculation methods: Lin, Resnik, Schlicker, and Wang; two model organisms: zebrafish and mouse; two evaluation curve types: ROC and precision-recall curves), the integrated networks outperform PPI networks and are better for predicting candidate genes for anatomical entities. Furthermore, the integrated networks prove better than both anatomy-based and PPI networks when the same anatomical profiles are not used for the evaluation. Together, these results indicate that the integration of the experimental knowledge *via* anatomy ontology increases the quality of the PPI networks, therefore, improving their candidate gene prediction accuracy.

There is a lack of developed bioinformatics tools and codes to address large-scale network integration problems. Moreover, built-in codes and libraries for semantic similarity calculations are not readily available for the Python programming language [53]. Therefore, the Python scripts written for this research will be extended to a Python library focused on large-scale network integration and construction of semantic networks using semantic similarity calculation methods.

An interesting future challenge would be to include quality terms using Phenotype and Trait Ontology (PATO) to analyze the genes associated with certain qualities of anatomical entities, such as the size or the presence and absence of an anatomical term [54]. For this task, a computational framework must be established to include composite entity-quality terms. Alternatively, phenotype ontologies, which already include quality of an anatomical entity, such as the Human Phenotype Ontology [55] and the Mammalian Phenotype Ontology [56], can be directly used for the integrative framework. However, based on Manda, et al. [57], incorporation of entity-quality terms may not significantly increase the amount of semantic information captured from phenotype profiles. Nonetheless, incorporation of additional phenotype information into gene networks, which has not been tested, may increase the performance and is a potential method to improve the network performance.

This integrative approach will be extremely useful for studying disease phenotypes in humans and other model organisms. Predicting disease genes using biological networks is extremely widespread due to the associated medical implications [15, 16, 58, 59], and the challenge is to improve the accuracy of the predictions. Using this integrative method, experimental knowledge regarding known gene-disease associations can be integrated with the human gene/PPI networks. For this purpose, Human Disease Ontology [60, 61] can be used instead the Uberon to semantically capture the gene-disease annotations. Therefore, the integrative framework used in this work is adaptable to a broad number of research questions and is a powerful tool for the bioinformatics community.

### Abbreviations

PPI: Protein-protein interaction
HMS-PCI: High-throughput mass-spectrometric protein complex identification
GO: Gene ontology
GO-BP: Gene Ontology-Biological Process
IC: Information content
ROC: Receiver operating characteristic
AUC: Area under the curve
PATO: Phenotype and Trait Ontology

## Acknowledgements

The authors thank J. P. Balhoff for assisting with the retrieval of gene-anatomical entity relationships from the Monarch Initiative repository. The semantic network construction scripts were implemented on the High Performance Computing clusters at the University of South Dakota and we are thankful to the HPC team for their assistance. The authors thank T. J. Vision, D. Goodman, B. Wone, W. M. Dahdul, and L. M. Jackson for their assistance and helpful comments to improve this research.

## Authors’ contributions

All authors planned and designed the experiments. PCF wrote the Python scripts for the analysis and performed the experiments under the supervision of PMM and EZ. All authors analyzed the results, read and approved the final manuscript.

## Availability of data and materials

The network files and the anatomy profiles used for the candidate gene predictions are available at https://doi.org/10.6084/m9.figshare.9973703.v2 and the Python scripts used for this analysis is available at https://doi.org/10.5281/zenodo.3470875.

## Ethics approval and consent to participate

Not applicable.

## Consent for publication

Not applicable.

